# First complete genome sequences of Zika virus from Midwest Brazil, 2015

**DOI:** 10.1101/521666

**Authors:** Carla Julia da Silva Pessoa Vieira, Laís Ceschini Machado, Lindomar José Pena, Roberta Vieira de Morais Bronzoni, Gabriel da Luz Wallau

## Abstract

Zika virus (ZIKV) has been intensively studied in Brazil since 2015-2016 epidemics, but little is known about the virus in Midwest region of the country. We report here two ZIKV complete genomes, which were isolated during arboviral surveillance in Sinop city, southern border of the Amazonian forest, Midwest Brazil, 2015.

---

Zika virus (ZIKV) is a mosquito-borne arbovirus that has a positive-sense single-stranded RNA genome. This virus belongs to the *Flaviviridae* family, *Flavivirus* genus, and has recently emerged as one of the most serious global public health threat. Symptoms are usually mild and up to 80% of ZIKV infections can be asymptomatic (*1*). However, ZIKV has been associated to neurological complications such as congenital Zika syndrome in newborns, Guillain-Barré syndrome and other neurological complications in adults (*2, 3, 4*).

ZIKV has been intensively studied since the 2015 epidemics in South America following the virus global spread (*5, 6, 7*). Although the study of full-length viral genomic sequences of clinical isolates can lead to a better understanding of molecular epidemiology, phylogeny and viral virulence, currently there are few available ZIKV complete sequences especially from regions of difficult access in Brazil. We report here a genomic investigation and phylogenetic study based on complete ZIKV genomes from patients living in Sinop, a city located 503 km north of the capital Cuiabá, in the State of Mato Grosso, Midwest region of Brazil.

We investigated 63 serum samples from patients presenting clinical symptoms of dengue-like infection, during an arboviral surveillance, from December 2015 to February 2016, in Sinop. These samples were tested for flaviviruses and alphaviruses by Multiplex-Nested RT-PCR (online Technical Appendix). Samples collection and research use was approved by the Ethics Committee from Júlio Müller University Hospital – UFMT (288.172/2013).

A total of six samples were positive for ZIKV by Nested-PCR. Two PCR-positive samples (BR/Sinop/H355/2015 and BR/Sinop/H366/2015) were amplified by real-time PCR, using primers, probe and PCR conditions as described (*8*) and sequenced on the MiSeq (Illumina) platform at the Technological Platform Core (online Technical Appendix). One sample (BR/Sinop/H355/2015) belonged to a 41-year-old female patient presenting petechial rash, back pain and arthralgia, and normal leukocyte (4,530/mm^3^) and platelet count (314,000/mm^3^). The other sample (BR/Sinop/H366/2015) was from a 30-year-old pregnant patient presenting fever, petechial rash, itch, diffuse myalgia and arthralgia, and normal leukocyte (6,390/mm^3^) and platelet count (140,000/mm^3^).

The entire ZIKV genomes from these samples were obtained with average coverage depth ranging from 280 to 9282 (Technical Appendix Table). Phylogenetic reconstruction of the obtained genomes showed that strain isolated from Sinop patients clustered within the Asian lineage (Figure, panel A) along with several other draft ZIKV genomes from the 2015-2016 epidemics. More specifically, one complete genome sequenced here clustered in a clade along with several other genomes from the North and Southeast Brazilian regions and genomes from Central America (355 sample - Figure, panel B, clade A) and the 366 genome clustered within another clade along with samples from Northeast and Southeast Brazilian regions as well as genomes from Central America (Figure, panel B, clade B). In addition, we could observe that the most closely related ZIKV genomes were detected in patients from the states of Tocantins, east border state of Mato Grosso, and from Paraíba state, located in the Northeast region, an area highly affected by the ZIKV epidemics (Figure, panel C).

We also characterized a number of nonsynonymous nucleotide mutations in the genomes obtained. We detected the previously reported amino acid mutations in one structural and four of the nonstructural proteins, including the A983V mutation NS1, which has been implicated in immune evasion and is conserved in all Asian strains detected after the French Polynesia outbreak (*9*) and a pre-M mutation (S139N) which has been associated with microcephaly in mice (*10*). In addition, we also found mutations specific for our samples: a NS2B mutation (I1398V), a NS3 mutation (T2068M) and a NS4B mutation (S2456L) (Technical Appendix Table).

**Technical Appendix Table:** Number of reads, quality and coverage for the genomes reported in this study. Genome coverage (%) was calculated relative to sequence KX197192 (10807 nt long).

| Isolate ID | Genbank<br>accession code | No. total<br>reads | No. mapped reads<br>(%) | Depth of<br>coverage | Genome<br>length (nt) | Amino acid<br>mutations and<br>protein location* |
| --- | --- | --- | --- | --- | --- | --- |
| H355 | MH513598 | 676,945 | 40,478 (5.98%) | 280.92 | 10640 | M1143V NS1 |
|  |  |  |  |  |  | I1398V NS2B |
|  |  |  |  |  |  | T2068M NS3 |
|  |  |  |  |  |  | S2456L NS4B |
| H366 | MH513599 | 1,454,943 | 1,337,597 (91.93%) | 9,282.85 | 10658 | M1143V NS1 |

Depth of coverage = Number of mapped reads x 75 (read length) / reference genome length (10,807).
\*Positions considering the reference genome polyprotein

The largest incidence of Zika in the Mato Grosso state occurred in 2015 (*11*). From 2015 to 2018, 6920 cases of DENV, 1392 of ZIKV, 12 of CHIKV were reported in the city (*12*). This report supports that the ZIKV circulating in this remote region of Brazil belongs to the Asian genotype and more specifically to two different clades found in other Brazilian states. Moreover, the close clustering with ZIKV sequences available from other Brazilian states which are far apart from Mato Grosso state highlights the rapid expansion of these strain between distant Brazilian regions. Lastly, we found two of ZIKV amino acid mutations that were associated with the virus pathogenicity and conserved in all ZIKV genomes obtained after 2014 and several other amino acid changes which their role remains unknown. Such findings highlight the need of rapid and efficient deployment of molecular surveillance in affected areas of difficult access (*5*).

**Legend Figure.**
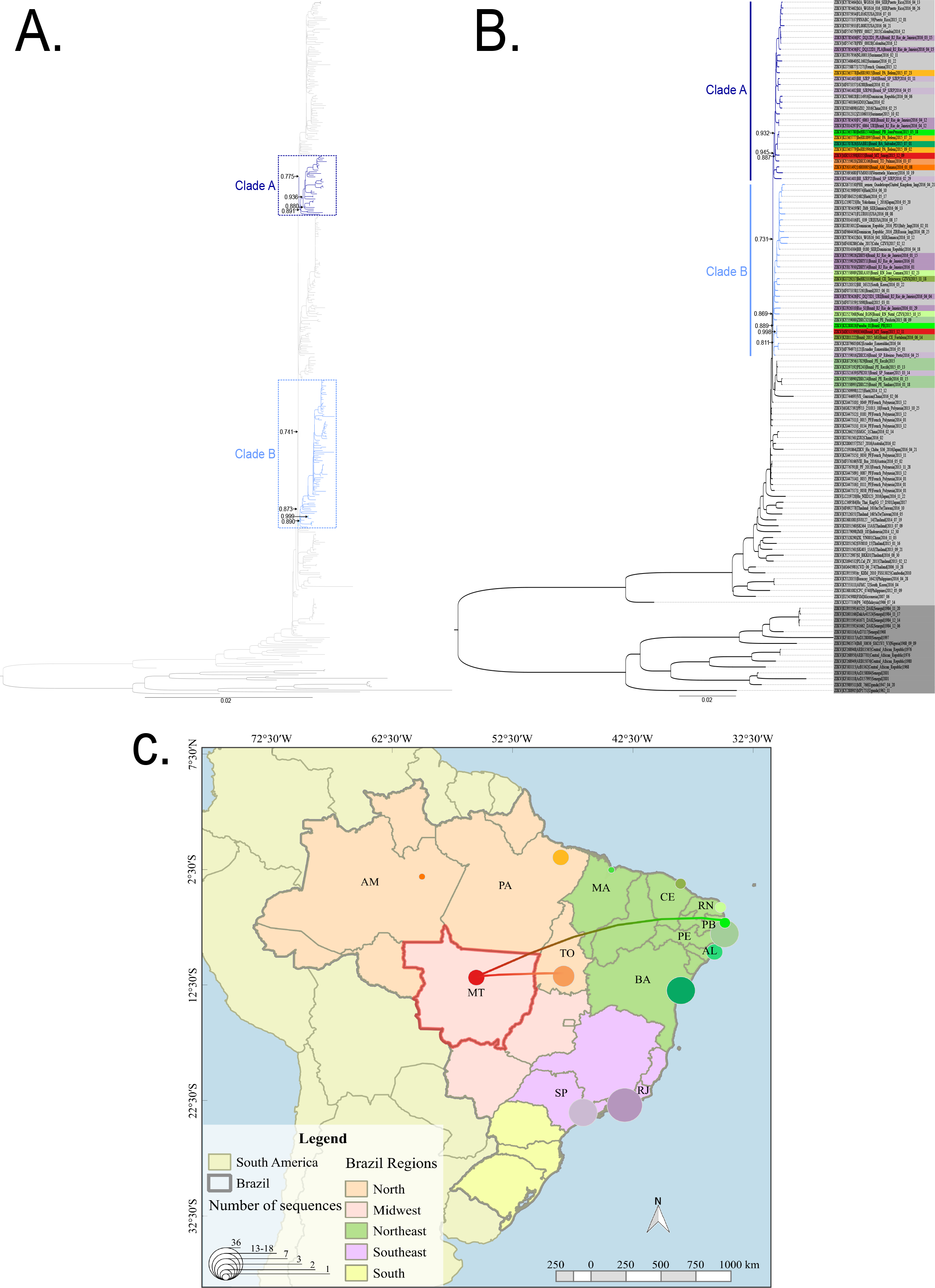
Phylogenetic reconstruction and positioning of ZIKV genomes obtained in this study using maximum likelihood. A - Full phylogeny including all available ZIKV complete and draft genomes > 9kb. B - Phylogenetic analysis of a subset of ZIKV sequences from the full ZIKV tree focusing on clades were the two ZIKV genomes obtained in this study clustered. C - Distribution of the number of ZIKV genomes sequenced per state and colored edges showing the possible route of ZIKV spread reaching Sinop based on the most closely related ZIKV genome from the previous phylogenetic analysis. Tip colors in panel B follow state colors in panel C. aLRT SH-like branch support of key clades are depicted with arrows in panels A and B.

## Technical Appendix

## Laboratory testing

Total RNA was extracted from 140 μL of patients’ sera using QIAmp viral RNA kit, followed by a reverse transcription using random primer, and Multiplex-Nested-RT-PCR using specific primers targeting NS5 and nsP1 regions, respectively (*1*). Briefly, we designed a new primer for ZIKV (CACTGGCCTCCTAGGCCCGTCCAT) and add it to the Multiplex-Nested-PCR for flaviviruses, along with DENV 1-4. We add CHIKV primer (TAGAGCAGGAAATTGATCCC) previously designed (*2*) to the Multiplex-Nested-PCR for alphaviruses.

## Genome sequencing

We used primers designed by the ZIBRA project (*3*) following as described (*4*) on the cDNA generated from the total RNA extracted directly from each sample. Samples were sequenced on the MiSeq (Illumina) platform at the Technological Platform Core at the Aggeu Magalhães Institute (IAM). Trimming and quality filtering of the recovered reads were performed with Trimmomatic v 0.36 (*5*) and quality checked with FastQC (http://www.bioinformatics.babra-ham.ac.uk/projects/fastqc/). These were subsequently mapped against the ZIKV BRPE243/2015 reference genome (KX197192) (*6*) using Bowtie 2 (*7*). The obtained sequences of the ZIKV genome have been deposited in GenBank with the accession number: MH513598 and MH513599.

## Phylogenetic Analyses

Multiple sequence alignment was performed by using MAFFT version 7 (http://mafft.cbrc.jp/alignment/software/); maximum-likelihood (ML) was determined by using PhyML version 3.0 (http://www.atgc-montpellier.fr/phyml/). Coding regions corresponding to the complete genomes from Sinop were aligned with all published and available near-complete Zika virus genomes (>9,000 nucleotides), belonging to the African and Asian genotype, totalizing 408 genomes, collected on GenBank. The ML phylogeny was reconstructed by using the best-fit general time-reversible (GTR) model with 4 gamma substitution rate categories (+ G) distributed with invariant sites (+ I) (GTR+G+I); for the tree search operation, we used SPR and statistical support for phylogenetic nodes was assessed by using aLRT SH-like. We also used FigTree v1.4.2 (*8*) for visualization and figure generation.

